# Nuku, a family of primate retrogenes derived from *KU70*

**DOI:** 10.1101/2020.12.02.408492

**Authors:** Paul A. Rowley, Aisha Ellahi, Kyudong Han, Jagdish Suresh Patel, Sara L. Sawyer

## Abstract

The ubiquitous DNA repair protein, Ku70p, has undergone extensive copy number expansion during primate evolution. Gene duplications of *KU70* have the hallmark of long interspersed element-1 (LINE-1) mediated retrotransposition with evidence of target-site duplications, the poly-A tails, and the absence of introns. Evolutionary analysis of this expanded family of *KU70*-derived “*NUKU*” retrogenes reveals that these genes are both ancient and also actively being created in extant primate species. *NUKU* retrogenes show evidence of functional divergence away from *KU70*, as evinced by their altered pattern of tissue expression and possible translation in the human testes. Molecular modeling predicted that mutations in Nuku2p at the interaction interface with Ku80p would prevent the assembly of the Ku heterodimer. The lack of Nuku2p-Ku80p interaction was confirmed by yeast two-hybrid assay, which contrasts the robust interaction of Ku70p-Ku80p. While several *NUKU* retrogenes appear to have been degraded by mutation, *NUKU2* shows evidence of positive natural selection, suggesting that this retrogene is undergoing neofunctionalization. Although Nuku proteins do not appear to antagonize retroviruses in cell culture, the observed expansion and rapid evolution of *NUKUs* could be being driven by alternative selective pressures related to infectious disease or an undefined role in primate physiology.

## INTRODUCTION

Protecting the integrity of a cell’s genetic material is important for both survival as well as for ensuring the faithful transmission of genes to daughter cells. Thus, DNA repair genes are conserved throughout the evolutionary history of prokaryotes and eukaryotes, with homologs present in every major organismal clade. A prime example is the *KU70* gene, involved in DNA double-strand break repair mediated by non-homologous end-joining (NHEJ). Human Ku70p and Ku80p together form the Ku heterodimer, a well-established initiator of the NHEJ pathway for DNA double-strand break repair [1–4]. In addition to its well-documented role in the NHEJ pathway, Ku70p is also involved in V(D)J recombination [5,6], telomere maintenance [7,8], Bax-mediated apoptosis [9], innate immune signaling [10–12], and is even involved in cell-cell adhesion and extracellular matrix remodeling at the cell membrane [13–15]. The *KU70* and *KU80* genes are present in eukaryotic and archaeal genomes, while in bacteria the role of the heterodimer is performed by a homodimer of the protein Ku [16,17].

Gene duplication is an important mechanism by which new genes arise. After gene duplication, multiple possible fates await the new gene copy, depending on the selective forces at play: decay, purifying selection, subfunctionalization, or neofunctionalization [18,19]. Retrogenes (previously known as ‘processed pseudogenes’) are a type of gene duplication created when retrotransposons erroneously reverse transcribe a cellular mRNA and insert the cDNA copy of the gene back into the host genome [20]. As a result, retrogenes often lack introns [21–23]. In addition, they can also be flanked by target-site duplications (TSDs), as is the case for mammalian LINE-1 mediated retrotransposition [24,25]. Retrotransposition and the subsequent formation of retrogenes is cited as having had a singular effect on primate and human evolution, with a so-called “burst” in retrogene formation during the last 63 million years having contributed to the emergence of many novel genes [26,27]. Approximately 3,771-18,700 retrocopies of human genes exist in the human genome, with about 10% of these found to express mRNA transcripts [28–30].

The main *KU70*-related gene duplication that is known is the ancient duplication that gave rise to *KU70* and *KU80*, and thereby the eukaryotic Ku heterodimer. Here, we report the description of five *KU70* retrogenes in the human genome, which we have named *NUKU1* – *NUKU5*. Four of these retrogenes are present in all simian primate genomes, and therefore predate the split between Old World monkeys and New World monkeys over 30 million years ago. However, a newer retrogene found on the human X chromosome, *NUKU5*, is specific to apes (human, gorilla, chimpanzee, and orangutan). *KU70* has spawned an unusual number of retrogene copies, as it is the only one out of 66 genes linked to DNA double-strand break repair to have five retrogenes in the human genome. While the original open reading frames appear to be disabled, there is evidence for expression of *NUKU2, NUKU4*, and *NUKU5* and a spliced transcript that exists for *NUKU2. NUKU2* has also evolved under positive selection, and functional tests of *NUKU* genes and molecular modeling simulations reveal that it has functionally diverged from *KU70* in two ways. First, whereas *KU70* is expressed in all tissues, *NUKU2, NUKU4*, and *NUKU5* display a tissue-specific expression pattern. Second, while Ku70p interacts with Ku80p, Nuku2p does not. Given the extensive functional characterizations of human *KU70* and *KU80* that have occurred over decades, it will now be of great interest to determine what potential role these additional Ku70-like proteins play in human biology.

## Results

### Five Ku70 Retrogenes in the Human Genome

Five open reading frames (ORFs) with high similarity to *KU70* were identified on four different human chromosomes (Figure 1A). Unlike the human *KU70* gene locus, each of the five copies lack introns. TSDs characteristic of LINE-1 mediated insertion were identified flanking each of the retrogenes, as were 3’ poly-A tails that are relics of the mRNA from which these genes arose (Figure 1A and 1B). All human retrogenes are between 89-97% identical to the parent *KU70* processed mRNA transcript and have been named *NUKU1* – *NUKU5*. Each of the five TSDs is unique, confirming that these copies represent five independent retrotransposition events, and did not arise from segmental duplication of an existing retrogene-containing region. Thus, the human genome contains one *KU70* gene and five LINE-1 mediated *NUKU* retrogenes.

**Figure 1.**
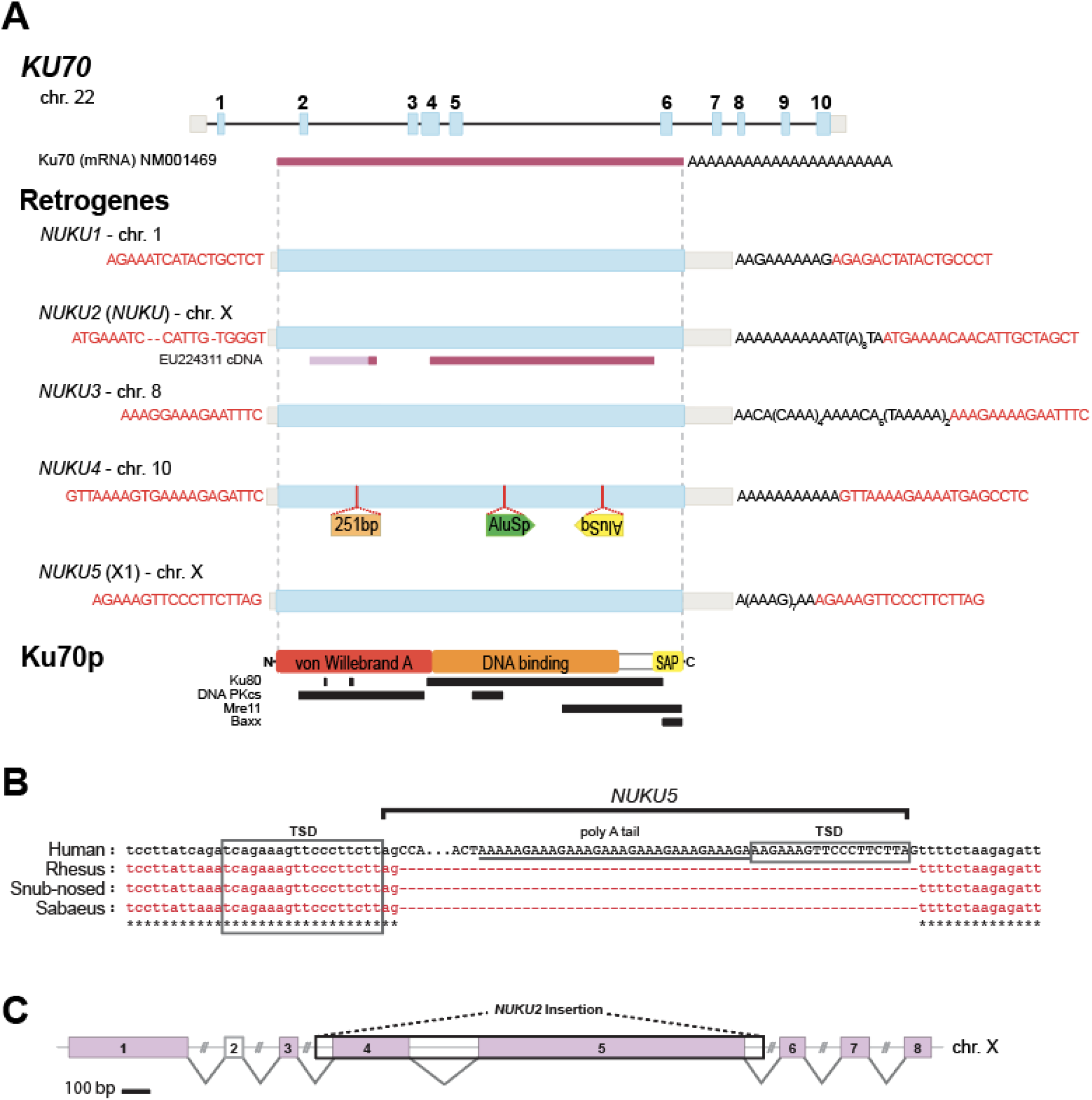
Identification of Five *KU70* Retrogenes in the Human Genome. A) A diagram of the *KU70* parent gene locus and the loci of its five retrogenes. Exons are shown in thick blue boxes and introns appear as black lines. 3’ and 5’ UTR structures are shown in light gray. Target-site duplication (TSD) sequences are highlighted in red text. B) Insertion of *NUKU5* in the human X chromosome compared to the syntenic locus of other Old World primates and evidence of LINE-1 mediated TSD. C) A lymphocyte-specific processed mRNA mapped to the human X chromosome with the insertion site of *NUKU2* boxed in black. Predicted splice sites are indicated between exons with 100% identity to the X chromosome (pink boxes). A significant match to exon 2 was not identified within the X chromosome.

We then analyzed several primate genomes for the presence of *KU70* retrogenes. Phylogenetic analysis (Figure 2A and S1) and inspection of pre-insertion target sites (as in Figure 1B and S2) defines the order in which these retrogenes arose, and places them at distinct positions in the tree of primate speciation (Figure 2B). These data show that four of the *KU70* retrogenes arose before the split between Old World and New World monkeys, over 30 million years ago (MYA), consistent with a burst of retrogene formation that has been reported in this time frame [26,27]. Remnants of *NUKU2* and *NUKU3* are present in the marmoset and squirrel monkey genomes (Figure 2A, S1), although they have experienced large subsequent deletions (Figure S2). We were unable to identify *NUKU1* in either the marmoset or squirrel monkey genomes (Figure S1) Comparing the syntenic location of *NUKU1* in both marmoset and squirrel monkeys to the human genome reveals large indels that prevents the reconstruction of the evolutionary history of the locus in New World monkeys (Figure S2). Since *NUKU1* is the most basally branching retrogene, we predict that it also predates the last common ancestor of the species being analyzed. Interestingly, the genomes of both marmoset and squirrel monkeys have acquired many additional *KU70* retrogenes that are not found in any of the other primate genomes investigated, meaning that these arose after the last common ancestor of New World and Old World monkeys (30-40 MYA) (Figure 2 and S1). The human genome contains one new retrogene, *NUKU5*, that is found in the genomes of chimpanzee and orangutan, but not in rhesus or marmoset. The pre-insertion site in the syntenic location in the rhesus macaque, snub-nosed monkey, and sabaeus monkey genomes are perfectly preserved (Figure 1B), confirming that this retrogene post-dates the split between Old World monkeys and hominoids that occurred approximately 20 MYA. Analysis of the genome of golden snub-nosed monkey also reveals the birth of a new *KU70* retrogene (*NUKU6*) with a TSD, remnants of a poly-A tail, which is absent from other Old World monkeys and humans (Figure S3). Thus, *KU70* retrogenes have been consistently birthed over a period lasting more than 30 million years, with evidence of continued retrogene birth in extant primate species.

**Figure 2.**
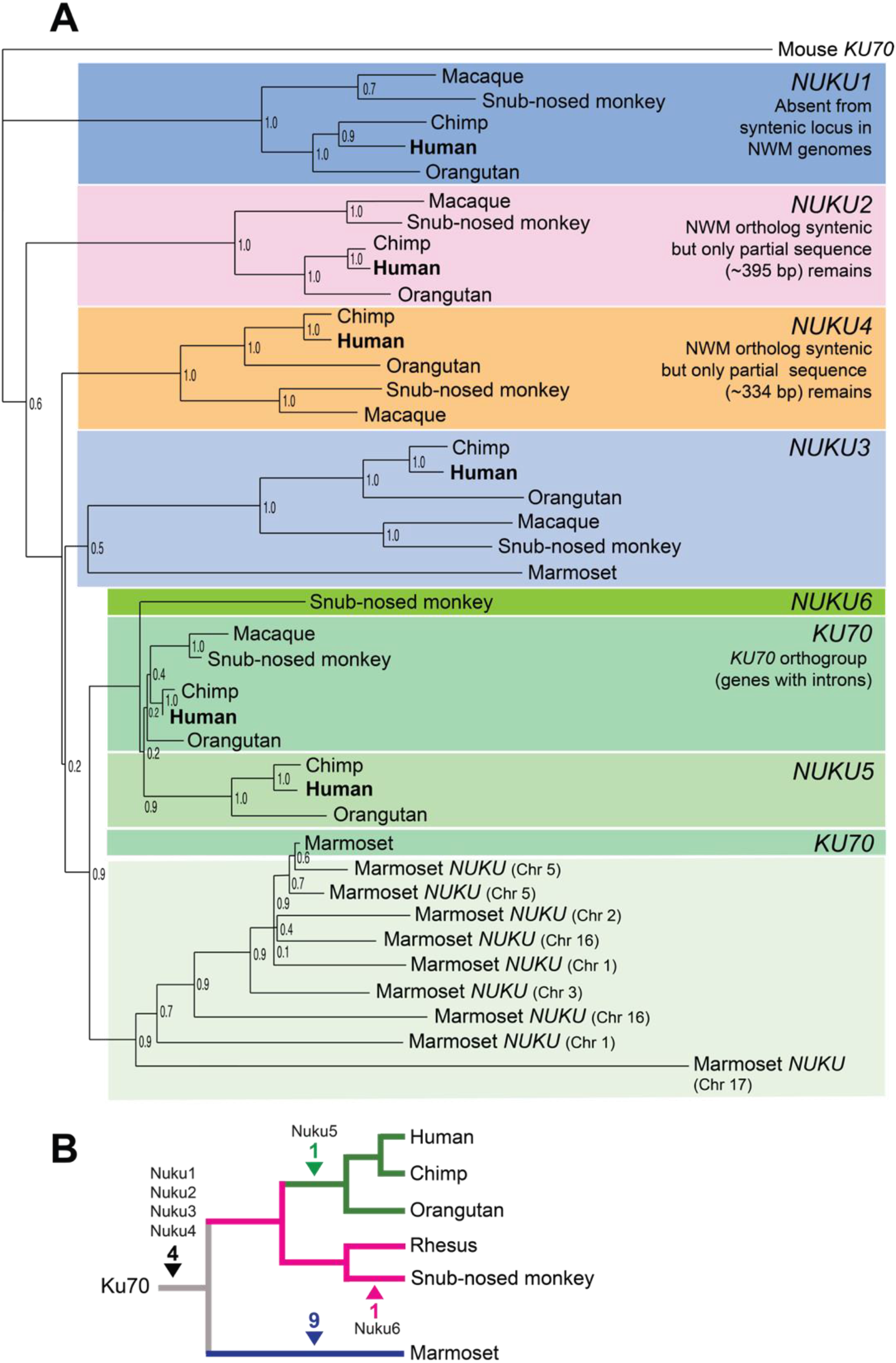
Phylogenetics and insertion sites of *KU70* derived retrogenes. A) Once the five *NUKU* retrogenes had been identified in the human genome, orthologous retrogenes were identified in other available primate genome projects through inspection of the syntenic target sites. A tree of these sequences is shown. Unless indicated, none of the genes on the tree contain introns. Bootstrap values generated with the maximum likelihood method are shown. Marmoset *NUKU3* was verified to be orthologous to the other *NUKU3* sequences by target site analysis. *NUKU2* and *NUKU4* are apparent in the marmoset genome, but are almost completely deleted, and therefore they were not included in the alignment used to make the tree. We were unable to locate the syntenic region of *NUKU1* in the Marmoset genome, indicating that this region may have been deleted (Figure S2). Marmoset-specific retrogenes were not named but are designated by the chromosome on which they are found. B) Based on the phylogenetic analysis and target site inspection, *NUKU1* – *NUKU4* predate the split between Old World monkeys, New World monkeys, and hominoids. *NUKU5* is specific to the great ape genomes analyzed and *NUKU6* is unique to the snub-nosed monkey. The marmoset genome has birthed 9 additional *KU70*-like retrogenes.

None of the ORFs in any of the primate *NUKU* retrogenes have been conserved in their full-length form as compared to *KU70*, and at first glance they all appear to be retropseudogenes. *NUKU3*, located on chromosome 10, has acquired two *Alu* insertions (*Alu*Sp and *Alu*Sq elements) and a 251 bp insertion of non-*KU70* related sequence in the middle of the coding region (Figure 1A). The ORF in *NUKU5* is approximately 75% the length of *KU70*, although *NUKU* ORFs are smaller, and the putative start codon of all of them is downstream of the *KU70* start codon. Surprisingly, a processed human mRNA transcript sequenced from lymphocytes (EU224311) was identified in the database that verifies the transcription and splicing of *NUKU2* on the X chromosome (Figure 1C). While we were unable to detect this spliced transcript by PCR, potentially because it is lymphocyte-specific, we performed 5’ and 3’ RACE to characterize the structure of a different unspliced transcript of *NUKU2* from total RNA isolated from the human testis (File S1). In conclusion, some of these human retrocopies express transcripts, including complex spliced transcripts.

### KU70 has an unusually large number of retrogenes

We were interested in determining whether the presence of five retrogenes of *KU70* in the human genome is typical for a gene involved in double-stranded break repair. Because some gene families might be more or less prone to retrogene formation and retention than others, we compared the number of retrogenes formed from *KU70* to other genes involved in DNA double-strand break repair. A list of all genes in the “double-strand break repair” biological process category (GO: 0006302) was compiled using the Gene Ontology (GO) database. Each was used as a query to identify retrogene copies elsewhere in the human genome. A retrogene was defined as any sequence match that 1) contains no introns, and 2) returns the parent gene when it itself is used to query the human genome (i.e. the gene and retrogene are reciprocal “best hits”). No criteria for conservation of the ORF was included, and some retrogenes appear to be degraded by mutation. In total, 51 double-strand break repair genes had no discernable retrogenes. Eleven genes (*MRE11, RAD21, FEN1, TRIP13, UBE2V2, PIR51, SHFM1, BRCC3, RNF168, OBFC2B*, and *RTEL1*) had one retrogene. Two genes, *SOD1* and *FAM175A*, had two retrogenes, and one gene, *UBE2N*, had four retrogenes. *KU70*, with five retrogene copies, is the only one out of 66 with five retrogene copies (Figure 3).

**Figure 3.**
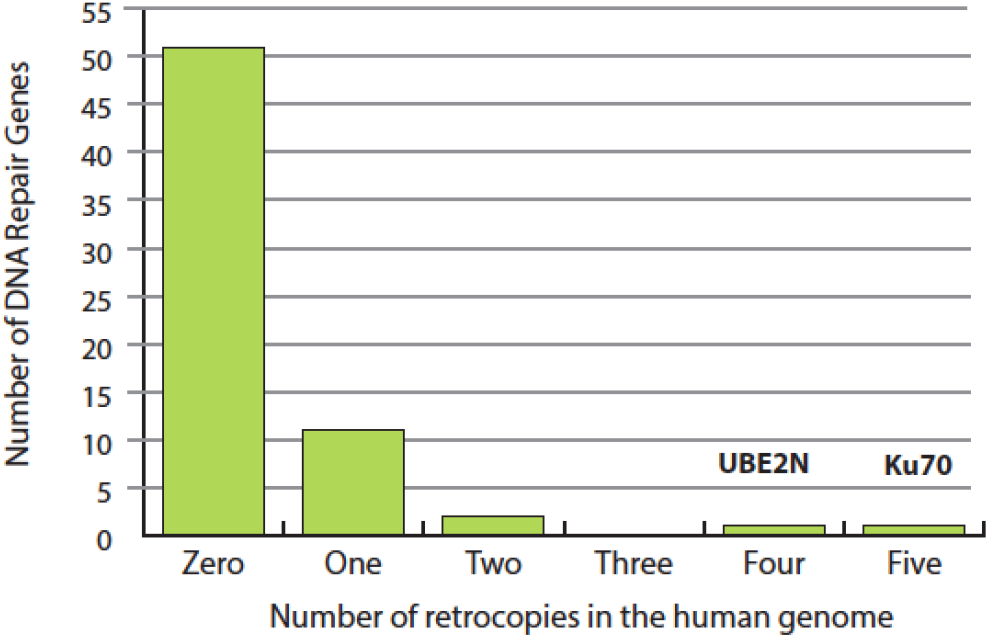
Prevalence of human retrogenes among double-strand break repair genes. The GO database was used to compile a list of 66 genes involved in DNA double-strand break repair. The human genome was searched for retrogene copies of each of these. The number if repair genes with 0, 1, 2, 3, 4, or 5 retrogene copies is shown. None of the 66 genes had more than 5 retrogene copies.

### NUKU2 has evolved under positive selection

There are three fates for any duplicated gene. A newly copied gene may be preserved by purifying selection if there is an adaptive advantage to having a second copy of the original gene. If the new gene copy is not expressed or confers no selective advantage, it will undergo neutral decay and accumulate point mutations and stop codons. Finally, if one of the duplicated genes is selected to evolve a novel function, this will occur through positive selection for advantageous mutations that arise and result in a period of relatively rapid sequence evolution in one of the copies. Each of these three fates can be read within the DNA sequence of duplicated genes after they have diverged. Looking at the evolutionary signatures recorded may offer clues as to the potential function of the retrogene and how it may relate to the parent gene’s function. Specifically, patterns of accumulation of non-synonymous versus synonymous mutational accumulation can be analyzed. Conserved genes like *KU70* would be expected to accumulate fewer non-synonymous changes than synonymous changes (dN/dS < 1). If a retrogene does not contribute to the fitness of the organism, it will accumulate these two types of changes at an equal rate (dN/dS = 1). However, if a retrogene acquires a new function and is selected for optimization of this function, it would bear the signature of increased non-synonymous mutation accumulation (dN/dS > 1).

The increased number of *NUKU* retrogenes is unexplained and could be rationalized if there is positive selection for their retention. The codeml program in the PAML package [31] was used to analyze the selective pressures that have acted on each of the *NUKU* retrogenes since they were formed. A tree of the human *KU70* and *NUKU* retrogenes was analyzed by the branch-sites model (Figure 4A). The analysis of patterns of non-synonymous and synonymous mutational accumulation can only be performed in ORFs, so a region at the C-terminal end of the retrogenes was analyzed because it is an ORF in all of the retrogenes except for *NUKU4*, which has experienced an *Alu* insertion in this region. The free-ratio model uses maximum likelihood to estimate a dN/dS ratio for each branch on the tree. As would be expected, the branch leading to *KU70* has a value of dN/dS = 0.45, indicating that non-synonymous changes have accumulated at a rate less than half of the rate of synonymous changes (Figure 4A). Three of the pseudogenes, *NUKU1, NUKU3*, and *NUKU5*, have a dN/dS signature not statistically different from 1, indicating neutral evolution of these genes. However, the branch along which *NUKU2* has been evolving shows a dN/dS value of 2.3. We retrieved the predicted ancestral sequence from the node marked “Anc,” which is the prediction of the *NUKU2* sequence as it looked at the time of retrotransposition (Figure 4A). Comparing this to the extant *NUKU2* sequence (Figure 4B) allowed us to determine that 17 non-synonymous mutations and three synonymous mutations have occurred in this region of the retrogene since it was formed more than 30 MYA. We used Monte Carlo simulation to determine that this rate of evolution is significantly greater than the neutral expectation of dN/dS = 1 (p = 0.007). The fact that at least one of these genes has evolved under positive selection agrees with the selected expansion of the *KU70* retrogene family that is observed (Figure 3).

**Figure 4.**
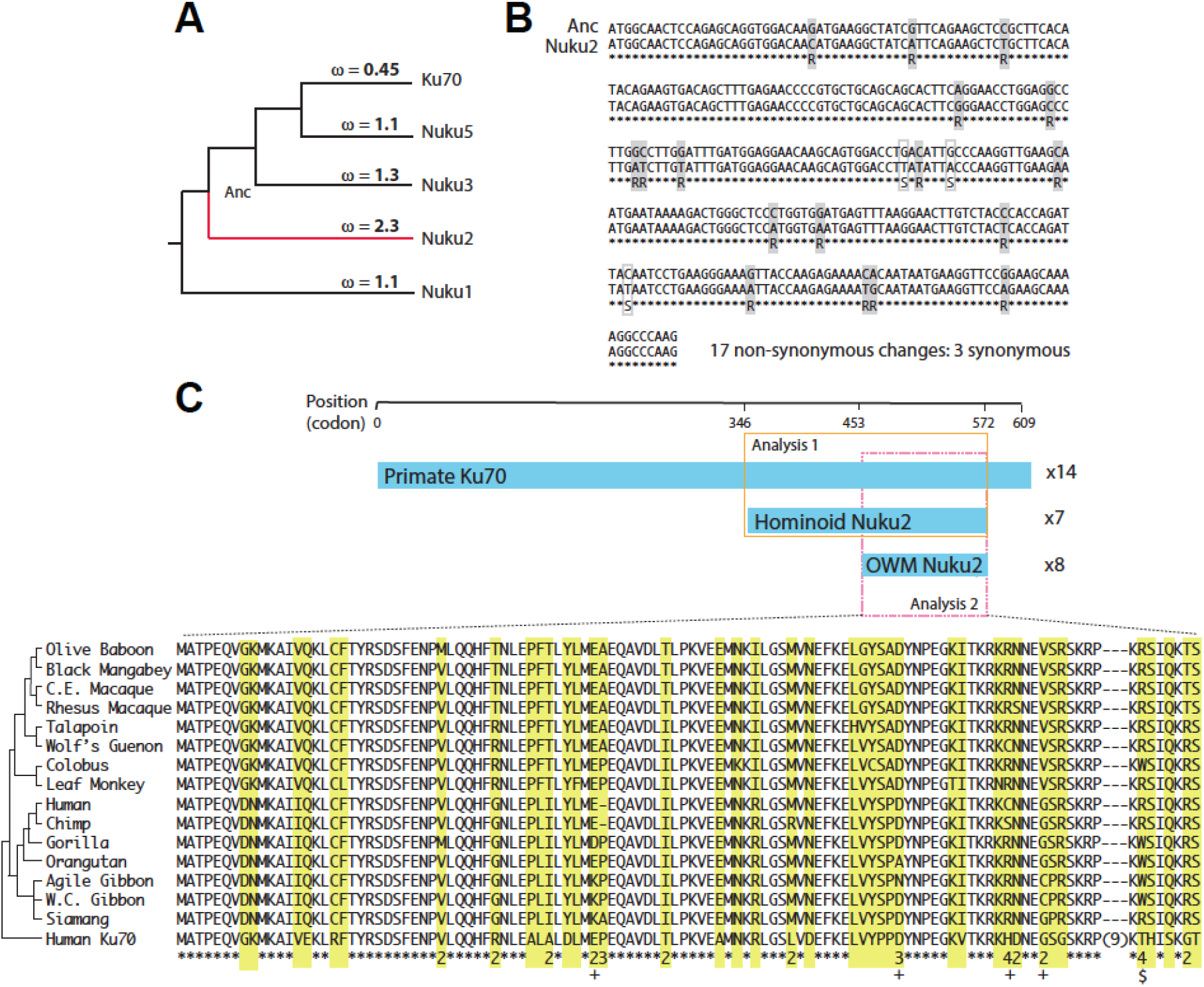
Molecular evolution of *KU70* retrogenes. A) Human *KU70* and four of the human *NUKU* retrogenes were aligned in the region of a common open reading frame. The branch-sites model assigned dN/dS values to each branch on the tree. These values summarize the evolution that has occurred since each retrogene was formed. “Anc” refers to the node representing the formation of *NUKU2*, and the predicted sequence at this node was generated by codeml. B) *NUKU2* is aligned to the “Anc” ancestral sequence in the region of the ORF which was analyzed in the analysis in panel A. Non-synonymous changes and synonymous changes are illustrated by gray and white boxes, respectively, in the alignment. C) *KU70* sequences were gathered for a total of 14 simian primate species, and *NUKU2* sequences were gathered from 15 species. All *NUKU2* sequences contain an ORF that is shorter than the *KU70* ORF, and it is even shorter in Old World monkeys than it is in hominoids. Two analyses of codon evolution were performed, one containing the sequences in the orange box (Analysis 1; longer ORF, *KU70* sequences plus 7 hominoid *NUKU2* sequence), and one containing the sequences in the pink box (Analysis 2; shorter ORF, *KU70* sequences plus all *NUKU2* sequences). The alignment shows the region that is an ORF in all genes. All *NUKU2* sequences are shown, with human *KU70* as an outgroup. In yellow are diverged sites, and numbers at the bottom indicate how many amino acid changes have occurred at those positions during *NUKU2* evolution (only indicated where dN/dS is greater than 1). The $ indicates a site that has changes from R to W three different times during *NUKU2* evolution. Plus signs indicate sites found to be under positive selection in the Analysis 1 branch-sites calculation (posterior probability > 0.5).

To further analyze the evolution of *NUKU2*, we determined the genetic sequence of *NUKU2* and *KU70* from 12 simian primate genomes (Table S1). Because it is expressed in all tissue types and contains multiple introns, *KU70* was amplified and sequenced from mRNA, whereas *NUKU2* was amplified and sequenced from genomic DNA. These sequences were combined with those available from several primate species with sequenced genome projects (human, chimpanzee, orangutan, and rhesus macaque), and genes were also re-sequenced from these species where appropriate. Our analysis includes only Old World monkey and hominoid species as *NUKU2* has been largely deleted in the marmoset and squirrel monkey genomes (Figure S2). Interestingly, the predicted ORF in human *NUKU2* (Figure 1A and 4C) was conserved in all hominoid species. In Old World monkeys, there was also a conserved ORF, but it was shorter due to an upstream stop codon leading to the potential use of an alternative ATG codon further downstream (Figure 4C). Since *NUKU* ORFs were predicted to be under positive selection and not *KU70*, we used the branch-sites model and specified all of the *NUKU2* branches as the foreground clade [32]. This allows us to look for positive selection of codons specifically in these species. Two analyses were performed, one with all *KU70* sequences and only the hominoid species where the longer reading frame was analyzed (orange box in Figure 4C), and one with all species where the shorter ORF was analyzed (pink box in Figure 4C). When the larger ORF was analyzed in hominoids only, it was estimated that that 9% of the codons in *NUKU2* had a dN/dS of 7.05. Comparison to the null model shows the inference of positive selection to be statistically significant (p = 0.029; Table S2). Support is not as strong when the shorter ORF in Old World monkey *NUKU2* was analyzed (p = 0.130), perhaps due to reduced statistical power.

**Table 1.**
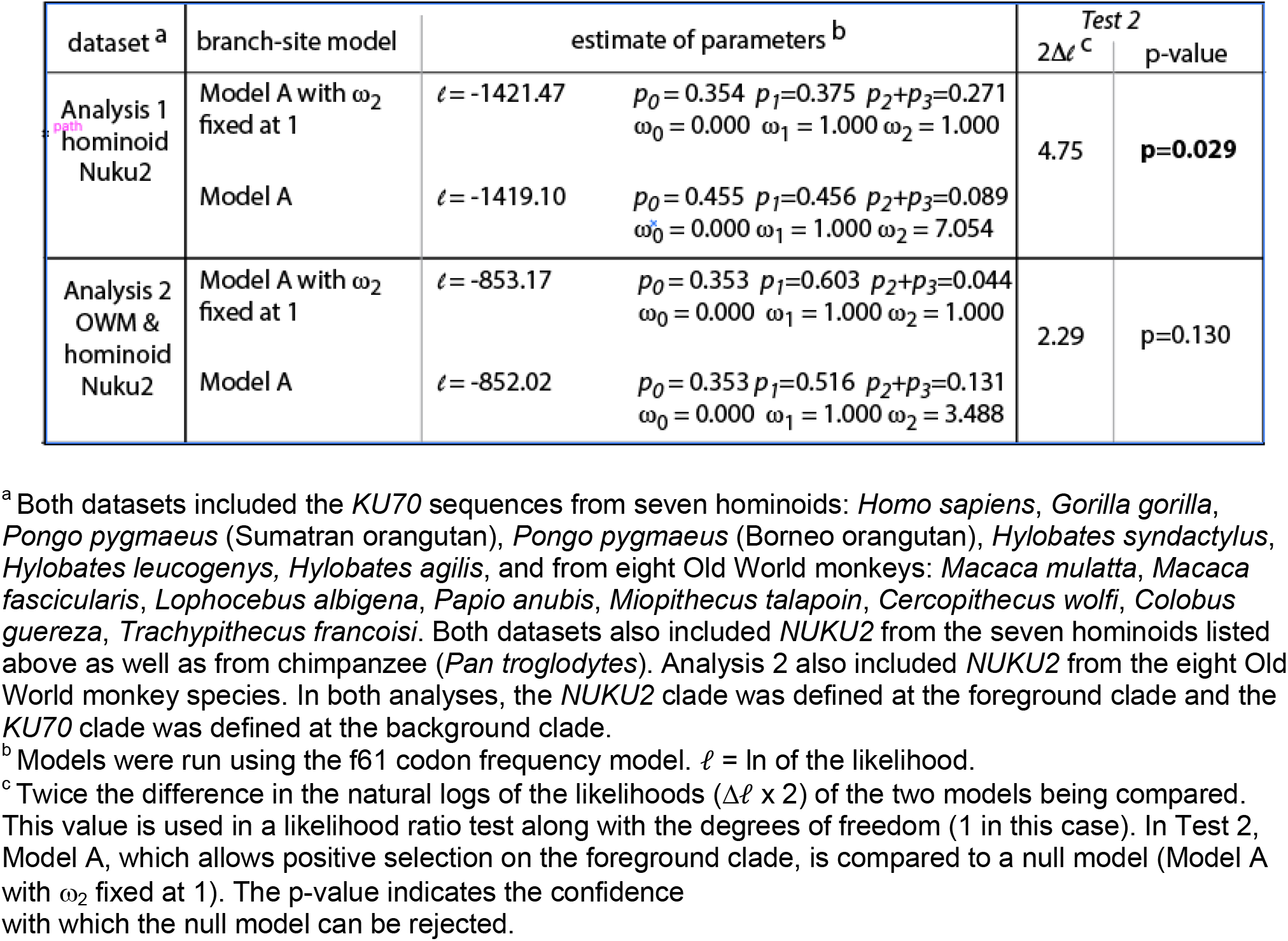
Brach-site for positive selection of Nuku2.

### NUKU2 has functionally diverged from Ku70

We designed PCR primers to specifically detect transcripts of the *NUKU2* retrogene. We used nested PCR with *NUKU2*-specific primers, determined the genetic sequence of all products and confirmed that they were a perfect match only to the *NUKU2* retrocopy. As shown, *NUKU2* is expressed in uterus, brain, testes, placenta, prostate, fetal liver, fetal brain, kidney, and spinal cord (Figure 5A). We confirmed the absence of contaminating genomic DNA by performing RT-PCR reactions in which the reverse transcriptase had been omitted. We also amplified *KU70* by a similar nested strategy, using primers located in two neighboring exons, to distinguish by size products of RT-PCR from PCR products that may be produced from contaminating genomic DNA. No genomic DNA was detected by this assay. This ubiquitous tissue expression pattern of *KU70* reflects its function as an essential housekeeping gene and is shown in other published datasets (Figure 5B) (GTex project version 7) [33]. We also found evidence for the tissue-specific expression of both *NUKU4* and *NUKU5* (Figure 5B). These results confirm that *NUKU2, NUKU4*, and *NUKU5* are expressed in humans, expression is tissue-specific, and tissue-specificity has diverged from that of *KU70*, likely due to new regulatory signals at their new genomic location.

**Figure 5.**
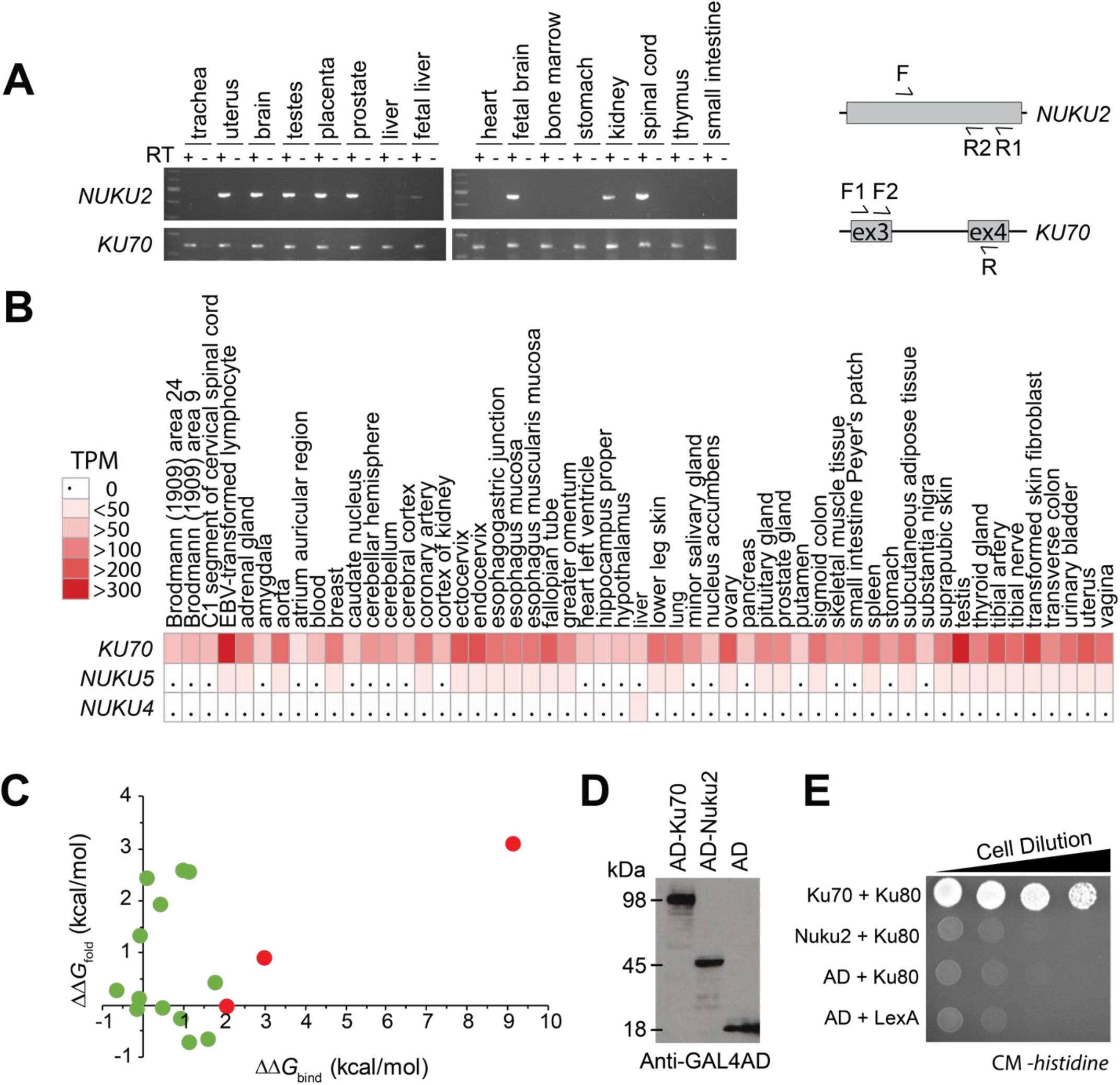
*NUKU* genes are functionally distinct from *KU70*. A) RT-PCR was used to analyze the expression of *NUKU2* and *KU70* from total mRNA harvested from different human tissues. Nested primer pairs are shown to the right. The product of a first-round RT-PCR reaction (primers F – R1) was then amplified with a second set of primers (F and R2), where R2 sits interior to R1. All three primers were designed to be specific to transcripts from *NUKU2*, as the ultimate base at the 3’ end of the primer placed such that it pairs with a base that is unique to *NUKU2* relative to the other five retrocopies. *NUKU2* does not have introns, but the *KU70* primers span an intron. Nested PCR with specific primers was also used to amplify the *KU70* transcript, which is different in size from the product obtained from genomic DNA. B) Relative tissue-specific expression patterns of *KU70, NUKU4*, and *NUKU5* measured in transcripts per million (TPM) [33]. C) For each Nuku2p mutation within 5 Å of Ku80p, ΔΔ*G*_bind_ was plotted on the x-axis, whereas ΔΔ*G*_fold_ was plotted on the y-axis. Mutations shown in green with x-axis values ΔΔ*G*_bind_ <2 kcal/mol and y-axis values -3 < ΔΔ*G*_fold_ < 3 kcal/mol are considered functional since they are likely to retain the ability to fold and bind. Mutations shown in red with x-axis values ΔΔ*G*_bind_ >2 kcal/mol and y-axis values -3 < ΔΔ*G*_fold_ < 3 kcal/mol are predicted to retain folding but disrupt Ku80p binding. D) Western blot confirming protein expression of each activation domain (AD) fusion construct in the yeast strains used for two-hybrid analysis. E) A yeast two-hybrid test assaying the interaction of Ku70p or Nuku2p with Ku80p. The Gal4 activation domain (AD) is either fused to Ku70p (top row), Nuku2p (second row), or expressed alone (third and fourth rows). LexA is a DNA binding domain and is either fused to Ku80p (top three rows) or expressed alone (bottom row). A positive interaction enables growth on complete media (CM) lacking histidine.

Ku70p is known to interact with Ku80p, thereby forming the Ku heterodimer that associates with broken ends of double-stranded DNA. To explore the potential biochemical function of a putative Nuku2 protein, compared to Ku70p, we examined the functional consequences of more than 10,000 mutational changes in Ku70p when bound to Ku80p using semi-empirical molecular modeling, as implemented by FoldX (Figure S4 and File S2) [34,35]. By comparing the amino acid changes between Ku70p and Nuku2p we individually modeled 27 non-synonymous mutations that are present in *NUKU2* onto the heterodimeric co-crystal of Ku70p-Ku80p (PDB:1JEY [36]) and measured the change in free energy for binding (ΔΔ*G*_bind_). The 11 mutations present in Nuku2p that were more than 5 Å from the Ku80p interface had an average ΔΔ*G*_bind_ of 0.04 kcal/mol (SD +/- 0.17), indicating that these changes would not be expected to disrupt Ku80p binding (Figure S4). The majority (81%) of the remaining 16 *NUKU2*-specific mutations that are within 5 Å of the Ku70p-Ku80p interface are also predicted to have little impact upon the interaction of these proteins (ΔΔ*G*_bind_ <2 kcal/mol; average 0.60 kcal/mol, SD +/- 0.74) (Figure 5C; green data points). However, four mutations at this interface (G349V, F410L, A494I, and T507I) had a ΔΔ*G*_bind_ >2 kcal/mol (Figure 5C; red data points). This indicates that these mutations alone would be predicted to disrupt the binding of Ku70p to Ku80p, and therefore, in combination are likely to prevent binding of Nuku2p to Ku80p. In addition, because Nuku2p is predicted to be truncated relative to Ku70p, there would be a 39% reduction in the surface area available for Ku80p binding from ∼9500 Å^2^ to ∼5800 Å^2^, which would also reduce the likelihood of a Nuku2p-Ku80p interaction (PISA analysis [37]). Analysis of disruptive mutations in hominoid *NUKU2* shows the presence of the same G349V, A494I, and T507I mutations that are found in human *NUKU2*. Only T507I appears within the *NUKU2* gene of Old World monkeys, in addition to a single disruptive mutation unique to colobus monkey (Y530C; ΔΔ*G*_bind_ >2) (Figure 4 and S4). Finally, non-synonymous mutations in *NUKU2* at sites under positive selection in primates have average ΔΔ*G*_bind_ and ΔΔ*G*_fold_ values of 0.33 (SD +/-0.45) and 1.00 (SD +/-1.46), respectively. This would suggest that these mutational changes were not driven by selection to disrupt Ku80p interaction or to alter Nuku2p folding.

Molecular modeling predicts that the truncation of *NUKU2* and several non-synonymous mutations disrupt an interaction with Ku80p. To validate these *in silico* predictions we used the yeast two-hybrid *in vivo* protein interaction assay to test the interaction of either Ku70p or Nuku2p with Ku80p. Ku70p and Nuku2p were both fused to the Gal4 activation domain (AD), and each construct was co-transformed with a plasmid encoding the LexA-Ku80p fusion protein (Figure 5D). Co-transformants of AD-Ku70p and LexA-Ku80p were able to grow on media lacking histidine, signifying a positive interaction. AD-Nuku2p and LexA-Ku80p were unable to interact and yeasts were unable to grow on histidine deficient plates. The LexA DNA binding-domain was also unable to interact with Ku80p or the AD (Figure 5E). Both the tissue-specific expression and inability to interact with Ku80p suggest that *NUKU2* has diverged from its parent gene *KU70* and potentially acquired new biological functions.

### Expression of NUKU2 and NUKU5 does not impact retrovirus replication

Ku is known to be important for the replication of many different viruses, including mammalian retroviruses and retrotransposons [38–42]. We considered that Nuku proteins could act to antagonize viral replication by mimicking Ku70p and evidence of positive selection might suggest host-virus antagonism (Figure 4). To test whether the expression of *NUKUs* might disrupt retroviral replication, we first confirmed the transient expression of *NUKU2* (human and rhesus macaque) and *NUKU5* (human) within the human HEK293T and HeLa cell lines (Figure S5). Twenty-four hours post-transfection these cell lines were transduced with GFP using VSV-G pseudotyped single-cycle human immunodeficiency virus 1 (HIV-1), feline immunodeficiency virus (FIV), and murine leukemia virus (MLV). Forty-eight hours post-infection the percentage GFP-expressing cells was measured using flow cytometry, and we found that *NUKU* expression did not affect retroviral transduction, relative to a control cell line expressing maltose binding protein (Figure S5).

### Detection of a Ku70-like protein encoded by a retrogene

Since several *NUKUs* appear to be transcriptionally active, we wished to address if either of these retrogenes was capable of producing a stable protein in human tissues. To do this, we needed to identify an antibody that would have cross-reactivity against these putative alternate protein forms of Ku70p. We assume that the two retrogenes most likely to be expressed as proteins are Nuku2p, for which we have documented tissue-specific expression, a spliced transcript, and positive selection, and Nuku5p, which is the youngest retrogene and the one with the longest ORF. We screened several anti-Ku70 polyclonal antibodies for cross reactivity with Ku70p, Nuku2p, and Nuku5p. We used Gal4AD-Ku70 or Gal4AD-Nuku fusion proteins expressed in yeast to test this, and we identified an antibody that specifically recognized all three constructs (Figure 6A). The protein band in HEK293T cell extracts shows the position of untagged Ku70p, and this antibody does not appear to cross-react with the endogenous copy of Ku70p in yeast. The tagged copy of human Ku70-AD is larger than the untagged version (Figure 6A, lane 2 versus lane 1). The tagged versions of Nuku2p and Nuku5p are shorter, due to the truncated ORFs in these two genes (Figure 1A).

**Figure 6.**
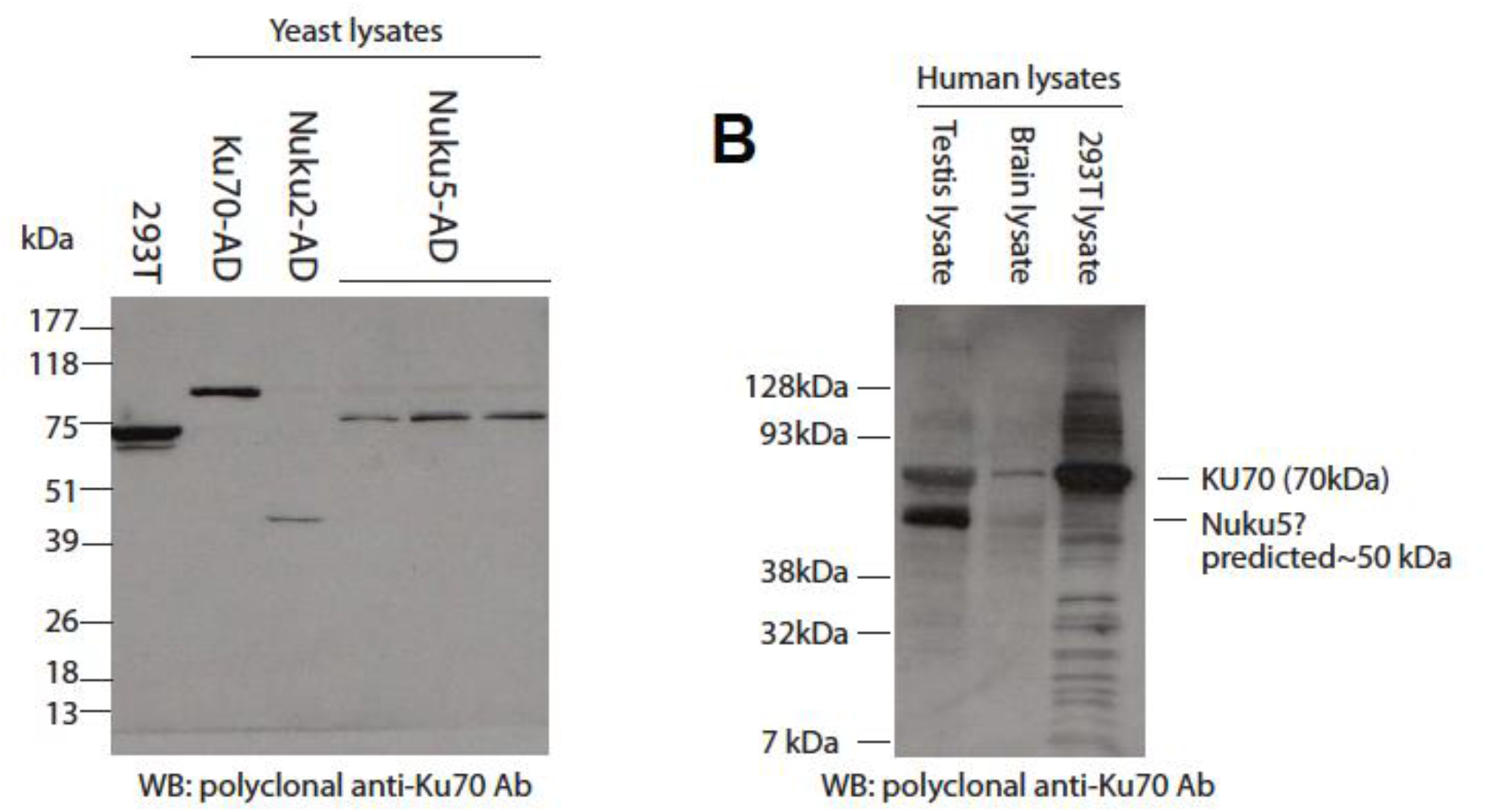
Detection of a putative *KU70* retrogene-encoded protein. A) Identification of an anti-Ku70 antibody that recognizes Nuku2p, Nuku5p and Ku70p. The ORFs of *KU70, NUKU2*, and *NUKU5* were fused to the *GAL4* activation domain (AD) and expressed in yeast. A Western blot of these proteins shows that a single polyclonal anti-Ku70 antibody recognizes all three Nuku fusion proteins (three independent transformants of Nuku5-AD are shown). B) Whole cell protein lysates from testis, brain, and HEK293T cells were purchased or cultured. The anti-Ku70 antibody characterized in panel A was used to probe these extracts.

We detected high levels of *NUKU2* transcription in brain and testis among other tissues (Figure 5B). Therefore, we probed protein lysates from human brain tissue, testis tissue, and from HEK293T cells with our anti-Ku70p antibody (Figure 6B). Protein lysates from HEK293T cells show only a single strong band at ∼70 kDa, the size of human Ku70p. This band is also evident in the testes and brain cell lysates. We did not detect a prominent band at ∼25 kDa in any of the samples, the predicted molecular weight of Nuku2p based on the transcript that we amplified by RACE (File S1). However, cell lysates from brain and testis tissues, but not HEK293T cells, show a second band at the predicted size of human Nuku5p (∼54 kDa), with the band being more prominent in testis than in brain.

## Discussion

*KU70* is highly conserved across primates, which contrasts other genes that are required for DNA repair that have been found to be evolving rapidly within humans and yeasts, potentially in response to selective pressure from viruses and retrotransposons [43–46]. Despite the conservation of *KU70*, we describe the accumulation and diversification of *KU70*-derived retrogenes within humans and non-human primates (*NUKUs*). The contribution of retrogenes to *de novo* gene formation and the evolution of novel gene functions has been extensively documented in different organisms [22,47–49]. *KU70* appears to be unique regarding the number of retrogenes that it has birthed relative to other genes required for NHEJ in primates. In addition to the expansion of the *NUKUs* we also have detected the rapid evolution and functional divergence of these retrogenes during primate speciation.

NHEJ is an important mechanism for DNA double-strand break repair in cellular organisms and is also important for the replication of DNA viruses and retroviruses/retrotransposons that generate DNA intermediates during their lifecycles. There are examples of NHEJ DNA repair mechanisms helping or hindering viral replication [50]. For example, lack of DNA-PK (DNA-dependent protein kinase holoenzyme, consisting of Ku70p, Ku80p, and DNA-PKcs) during HIV replication results in reduced viral integration and an increase in cellular apoptosis due to integrase-mediated DNA damage [39,42]. Also, loss of Ku70p causes the proteasome-mediated degradation of the viral integrase [38]. Retrotransposons and adenovirus have also been shown to be sensitive to the loss of Ku [40,41,51]. Bacteriophages encode Ku homologs that recruit other host DNA repair proteins and appear to protect phage DNA from degradation [52,53]. Furthermore, the hijack of NHEJ machinery is not specific to viruses as the bacterial pathogen *Rickettsia conorii* binds to cell surface-exposed Ku70p as its receptor for cell entry [13–15]. In these cases, it is apparent that NHEJ machinery (including Ku70p) is aiding the replication and survival of viruses and bacteria. Conversely, there are many examples of DNA viruses that encode protein effectors that actively disrupt the function of NHEJ. Specifically, adenoviruses prevent the concatenation of their genomes by NHEJ machinery by producing the proteins E4-34 kDa and E4-11 kDa that bind DNA-PK and inhibit NHEJ [54]. Human T-cell leukemia virus type-1 proteins Tax and HBZ and the agnoprotein of JC virus bind and interfere with the function of DNA-PK, impairing DNA repair and aiding cellular transformation [55–57]. Viral proteins also block the activity of DNA-PK as a pattern recognition receptor that binds cytoplasmic DNAs triggering innate immune signaling mechanisms mediated by IFN regulatory factor 3 (IRF-3), TANK-binding kinase 1 (TBK1), and stimulator of interferon genes (STING) [10–12]. DNA-PK has been shown to be directly targeted by the vaccinia virus effectors C4 and C16 by binding Ku and preventing interaction with DNAs and triggering of innate immune signaling pathways [58,59]. The abundance of viruses and bacteria that subvert the function of the DNA-PK suggests that the *NUKUs* could play a role as dominant-negative proteins that would bind viral effectors. It has already been shown in higher eukaryotes that a dominant negative Ku80p with an N-terminal extension (Ku80/Ku86-autoantigen-related protein-1 (KARP-1)) interferes with DNA-PKcs activity causing X-ray hypersensitivity when expressed in cell lines [60]. However, molecular modeling studies and empirical binding assays show that Nuku2p does not bind Ku80p and would therefore not be predicted to assemble as a component of DNA-PK. Furthermore, *NUKU2* appears to have only maintained coding capacity within the C-terminal domain, which is required for binding to DNA, Mre11p, and Bax, whereas the N-terminal domain binds to DNA-PKcs and Ku80p [9,61]. Therefore, we would expect that *NUKU2* would not influence DNA-PK function, V(D)J recombination, or telomere maintenance, but might still be competent as a transcription factor, or regulate apoptosis and NHEJ by binding Mre11p or Bax, respectively [9,65]. The observed expansion of *KU70* retrogenes and the rapid evolution of *NUKU2* could have been driven by evolutionary conflict with viruses or other pathogenic microorganisms free from the constrains of maintaining DNA repair of innate signaling functions. Indeed, the retrotransposition of genes involved in innate immunity can create new host restriction factors to fight rapidly evolving viruses [62–65]. Although we do not observe any significant effect of *NUKU* expression upon retrovirus infection in tissue culture, it remains plausible that other viruses know to interfere with DNA-PK or directly interact with Ku70p (i.e. JC virus agnoprotein or adenovirus E1A) might be sensitive to the presence of *NUKUs* [51,57]. Alternatively, as we have detected tissue-specific transcription from *NUKUs* it is also possible that they might have a function in the regulation of *KU70* expression as antisense transcripts [66]. Altogether, these data suggest that primate-specific *NUKUs* are significantly altered compared to *KU70* in their expression and protein-coding capacity. Our analyses suggest that their structure and function differ from *KU70* and that they have evolved rapidly during primate speciation. However, it remains to be further investigated the biological function of these retrogenes, which is complicated by the multifaceted role of Ku70 in the cell.

## Materials & Methods

### Identification & classification of retrogenes

The *KU70* coding sequence was used as a query in the UCSC genome browser against the human genome (http://genome.ucsc.edu/, March 2006 NCBI36/hg18 assembly). Six top hits of Blat scores were identified, the topmost of which matched with 100% sequence identity to the original *KU70* gene. The next five hits appeared as retrocopies upon closer inspection. *NUKU* orthologs from chimpanzee, orangutan, and rhesus macaque were also obtained using this method. For inspection of insertion sites in the marmoset genome, the calJac1 and calJac3 assemblies were used. All other insertion sites were interrogated using the current version of primate genomes found on the UCSC genome browser (https://genome.ucsc.edu). The phylogenetic trees of *KU70* and *NUKU* sequences were built with MEGA (maximum likelihood method).

The GO term “double-strand break repair” was queried in the GO database (GO term ID 0006302). Because not all genes have been fully annotated and assigned to appropriate GO categories (leading to exclusion of certain relevant genes from this list), we combined genes assigned to this GO category in either *Homo sapiens, Mus musculus*, and *Rat norvegicus*. This resulted in a list of 66 genes (Table S3). cDNA coding sequences for all 66 hits were retrieved from NCBI. In the case of genes with multiple transcript variants or splicing variants, the longest transcript was used. To find retrocopies of each gene, cDNA sequences were used as queries in the UCSC human genome database (hg18). Retrocopies were defined as hits in the human genome that met the following two criteria: 1) they lack introns (RepeatMasker was used to differentiate introns from transposable element insertions), and 2) they match the parent gene in a reciprocal best hit analysis of the human genome. Reciprocal best hit analysis was performed by taking each putative retrocopy and using the BLAST server at NCBI to query the human RefSeq mRNA database.

### Sequencing *KU70* and *NUKU* orthologs

*KU70* orthologs and *NUKU2* ORF orthologs were sequenced from mRNA-derived cDNA for Ku70 and from genomic DNA for *NUKU2* from 12 primates: gorilla (*Gorilla gorilla)*, agile gibbon (*Hylobates agilis*), colobus, crab-eating macaque (*Macaca fascicularis*), gibbon (*Pongidae Hylobates syndactylus*), leaf monkey, Borneo orangutan, talapoin, white-cheeked gibbon, olive baboon, black mangabey, and Wolf’s guenon. Genes were PCR-amplified using the strategy described in Table S4 and sequenced with primers shown in Table S5. The full structure of the *NUKU2* transcript was determined with 5’ and 3’ RACE using the GeneRacer kit (Invitrogen), and testicle total RNA (Ambion, catalog #7972). All nucleotide sequences are provided within File S3.

### Evolutionary analysis of *KU70* retrogenes

Sequences of the human *KU70/NUKU* paralogs were collected from the UCSC genome browser and aligned using ClustalX. Sequences were analyzed under the free-ratio model implemented in the codeml program of PAML 3.14. In order to determine whether dN/dS > 1 on the *NUKU2* branch, we made a pairwise comparison between the Anc sequence (generated by codeml) and *NUKU2*. K-estimator [67] was used to run Monte Carlo simulations of neutral evolution of these sequences, creating a null distribution from which a p-value could be derived.

The branch-site test allows identification of positive selection that might be limited to a subset of codons along only a subset of the branches being analyzed [32]. To implement this test, multiple alignments were fitted to the branch-sites models Model A (positive selection model, codon values of dN/dS along background branches are fit into two site classes, one (ω_0_) between 0 and 1 and one (ω_1_) equal to 1, on the foreground branches a third site class is allowed (ω_2_) with dN/dS > 1), and Model A with fixed ω_2_ = 1 (null model, similar to Model A except the foreground ω_2_ value is fixed at 1). *NUKU2* branches (back to their last common ancestor) were defined as the “foreground” clade, with all other branches in the tree being defined as background branches. The likelihood of Model A is compared to the likelihood of the null model with a likelihood ratio test.

### *NUKU* expression in human tissues

Total RNA from human tissues was purchased from Clontech (catalog number 636643). Most of these samples represent pooled RNA from multiple individuals (between 2 and 63 individuals). First-strand cDNA was produced with the NEB Protoscript II kit (E6400S), using a dT_23_ primer that anneals indiscriminately to poly-A tails on mRNA molecules. First-strand reactions were carried out twice in parallel for each tissue, one with reverse transcriptase (RT), and one with water added instead of RT (indicated by +/- RT on figure). First-strand cDNA was then amplified with *KU70*- and *NUKU*-specific primers using Invitrogen PCR Supermix HiFi (cat 10790020). In order to increase specificity, two successive PCRs were performed. In the first round of PCR, 20 cycles were performed using primers specific to that gene, along with 2 μL of first-strand cDNA as template. In the second cycle, 0.5 μL of the first round PCR reaction was used as template, and one of the gene-specific primers was substituted with a nested primer (F2 or R2 in diagram). In this round, amplification was performed for 40 cycles, and 2 μL of the final product was then run on a 2% agarose gel for separation. Primers used were: SS004 (Nuku F), SS011 (Nuku R1), SS009 (Nuku R2), SS030 (Ku70 F1), SS031 (Ku70 F2), and SS032 (Ku70 R) [ADD THESE TO PRIMER LIST]. The *KU70*-specific primers span an intron so that cDNA can be differentiated from the product that would be produced from genomic DNA. There are no introns in Nuku. Products were sequenced to confirm that they unambiguously represent *KU70* or *NUKU*.

### Molecular modeling of *NUKU2* using FoldX

To understand the effect of single missense variation on Ku70p stability (i.e. folding) and its binding with Ku80p, we estimated both folding and binding ΔΔ*G* values (difference of free energies between wild-type and the mutant) using FoldX software [34]. To run FoldX calculations, X-ray crystal structure of the human Ku heterodimer was first downloaded from Protein Data Bank (PDB id: 1JEQ) [36]. The file was modified to remove all but the two chains of Ku70p and Ku80p. There were several residues that were missing in both the chains of the protein complex. These missing residues were not modeled to complete the structure of the complex before running FoldX calculations for the following two reasons: 1) Missing residues were either at the terminal ends or in the disordered region hence they are difficult to build using the molecular modeling software and, 2) the gaps in the input X-ray structure does not affect the performance of the FoldX software as it relies on rotamer libraries to model any mutation at a particular site and semi-empirical scoring function to estimate ΔΔG values [35]. The clean starting structure of Ku70p-Ku80p complex was then used to create mutant models and subsequently estimate both binding ΔΔG and folding ΔΔG values. We started by performing 6 rounds of minimization of the protein complex using the RepairPDB command to obtain convergence of the potential energy. All 19 possible single amino acid mutations at each site on Ku70 (548 amino acid residues × 19 possible substitutions) were then generated using BuildModel. Finally, folding ΔΔ*G* values were estimated using Stability command on Ku70 and AnalyseComplex command was used to estimate the effect of each modeled mutation on Ku70p-Ku80p binding i.e. binding ΔΔ*G* values.

### Yeast two-hybrid assay

We used the LexA-Gal4 yeast two-hybrid system, which employs the LexA DNA-binding domain (DBD) and the Gal4-activation domain (Gal4-AD) with the yeast strain EAY1098 (*His3, Leu2, Trp1*, genotype). If the candidate proteins interact, the DNA-binding domain and activation domain will be in close proximity and will be able to drive the transcription of a *HIS3* reporter gene downstream of the LexA promoter. The Clontech pGADT7 plasmid, which creates an N-terminal fusion protein between a gene of interest and the Gal4 activation domain, was engineered to carry the full 1,830 bp coding sequence of human *KU70*. Another pGADT7 vector was engineered to carry the full 654 bp *NUKU2* open reading frame. The full-length coding sequence of human *KU80* (2,199 bp) was cloned into the LexA expression vector pBTM116, which creates an N-terminal fusion protein between the inserted gene and the LexA DNA binding domain. All cloning was done with TA-vectors and plasmids compatible with the Gateway system (Invitrogen). EAY1098 was transformed using the standard Lithium-acetate PEG transformation protocol with the following plasmid pairs: pGADT7-Ku70 and pLexA-Ku80; pGADT7-Nuku and pLexA-Ku80; pGADT7 and pLexA-Ku80; and pGADT7 and pLexA. Transformants were selected on leucine and tryptophan drop-out media to select for and stimulate expression of plasmids. After two days growth at 30°C, saturated cultures at an OD_600_ of 2.7-2.8 were diluted and plated onto media lacking histidine in addition to leucine and tryptophan to stimulate *HIS3* gene reporter expression. Growth was observed three days post-plating.

### Western blots

30 μg of denatured protein lysate was loaded onto 10% Tris-HCl polyacrylamide gels and then transferred onto a nitrocellulose membrane. Membrane was blocked overnight in 5% milk-TBS + 1% Tween and incubated the next day with a primary antibody directed against the Gal4-activation domain (1:5,000 dilution; Clontech, cat # 630402) or against human Ku70p (1:1,000 dilution; GeneTex, cat # GTX101820). The secondary antibody for Gal4 probes was goat anti-mouse-HRP (1:1,500; Fisher, cat #32430), and for Ku70p probes was goat anti-rabbit-HRP (1:1,500 dilution; Fisher cat. #32460). Signal was detected using ECL plus reagents (VWR cat #95040-056). For analysis of two-hybrid constructs, total protein from yeast strains prepared using the glass-bead disruption method. 50 mL yeast cultures were grown to OD_600_ 0.5-0.7 and were pelleted. This pellet was suspended in disruption buffer: 20 mM Tris-HCl, pH 7.9, 10 mM MgCl_2_, 1 mM EDTA, 5% glycerol, 0.3 M (NH_3_)SO_4_, with 1 mM DTT, 1 mM PMSF, and Protease inhibitor cocktail (Roche). Acid-washed glass beads were added and cells were vortexed for a total of 10 minutes.

### Western Blot analysis of Ku70 retrogenes

Human brain and testes tissue total protein lysates were purchased from ProSci Incorporated (catalogue numbers 1303 and 1313, respectively). HEK293T cells were grown in standard DMEM with 10% fetal bovine serum in 75 cm^3^ tissue culture flasks. Total protein was prepared using the reagents and protocol described in the Qiagen Mammalian Protein preparation kit. Protein was quantified using Pierce Coomassie Bradford Assay reagent. About 30 μg of protein was separated using polyacrylamide gel electrophoresis on a Tris-HCl gel and transferred to a nitrocellulose membrane. Membranes were blotted with 1:1000 dilution of the Ku70p antibody raised in rabbit (GeneTex XRCC6 antibody, Cat.# GTX101820). Secondary antibody of Goat anti-rabbit conjugated to horseradish peroxidase at 1:1500 dilution was used (Cat. #32460 Thermo Scientific Pierce Goat anti-Rabbit IgG, Peroxidase Conjugated). Maltose binding protein/hemagglutinin-tagged Nuku proteins were detected using an anti-HA peroxidase-conjugated monoclonal rat antibody (3F10; 12013819001 (Roche)).

### Virus infection assays

Human HEK293T (4 × 10^5^) and HeLa (4 × 10^4^) cells seeded in 12-well dishes (DMEM growth medium with 10% fetal bovine serum) and were grown at 37°C with 5% CO_2_ for 24 hours until reaching a confluency of ∼75%. Each well was transiently transfected with 800 μg of plasmid encoding either human *NUKU5, NUKU2* or rhesus macaque *NUKU2* in addition to a transfection control plasmid expressing RFP. After 24 hours incubation, each well was trypsinized and the HEK293T (2 × 10^5^) and HeLa (4 × 10^4^) cells used to seed three wells of a 24-well dish. After 24 hours of incubation at 37°C (5% CO_2_) monolayers with a confluency of ∼50% were infected with VSV-G pseudotyped HIV, FIV, or MLV containing a GFP reporter gene. After 48 hours, cells were trypsinized and fixed with 1% paraformaldehyde by incubating for 1 hour at 4°C. GFP and RFP positive transduced cells were detected by flow cytometry using appropriate compensation controls to account for spectral overlap of fluorophores.

### Data Availability Statement

Strains and plasmids are available upon request. The authors affirm that all data necessary for confirming the conclusions of the article are present within the article, figures, files, and tables.

## Acknowledgements

Research reported in this publication was supported by the National Institute Of General Medical Sciences of the National Institutes of Health under Award Number P20GM104420 (PAR and JSP). The content is solely the responsibility of the authors and does not necessarily represent the official views of the National Institutes of Health. SLS is a Burroughs Wellcome Fund Investigator in the Pathogenesis of Infectious Disease.

## Supplementary Figures

**Figure S1.**
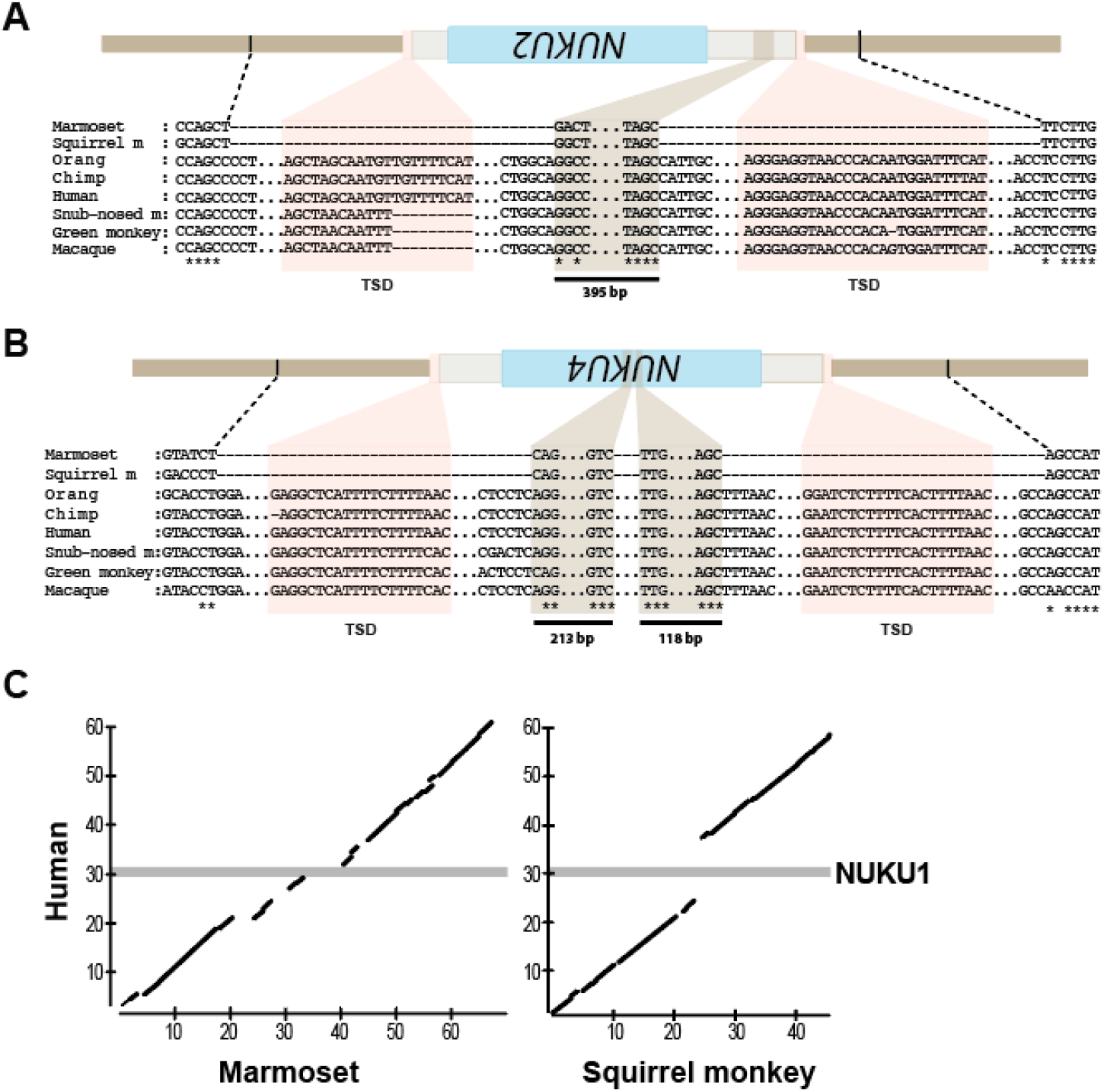
Evidence of the gain and loss of *NUKU* genes across different primate species. Remnants of (A) *NUKU2* and (B) *NUKU4* sequences in New World monkeys after gene deletion. Alignments show the sequence present at the syntenic position in each primate species. (C) Dot plot representation of the absence of *NUKU1* in marmoset and squirrel monkey genomes compared to the syntenic genome position in the human genome.

**Figure S2.**
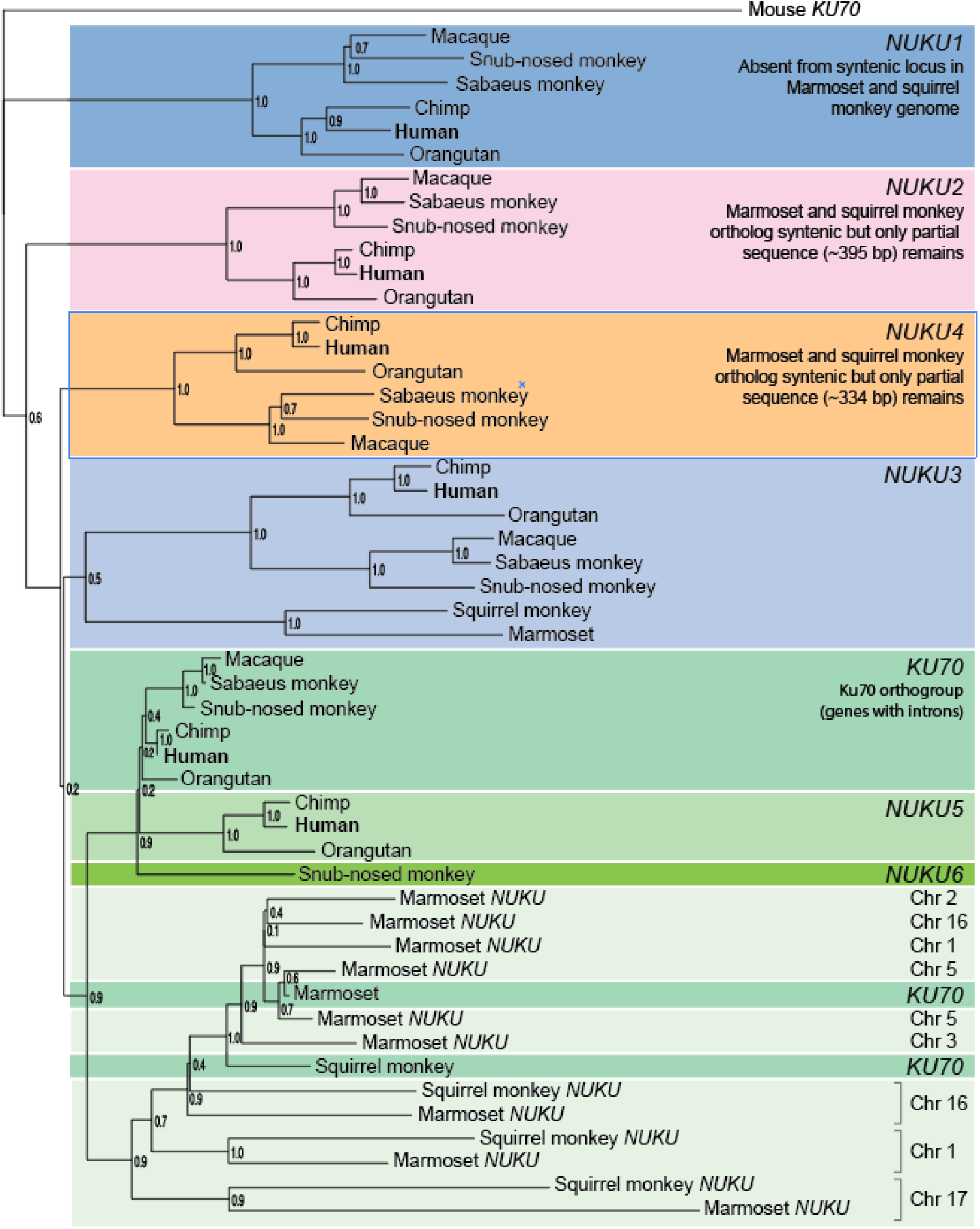
Phylogenetics of *KU70* derived retrogenes in eight primate species. A tree of *KU70* derived retrogenes sequences with Bootstrap values generated with the maximum likelihood method.

**Figure S3.**
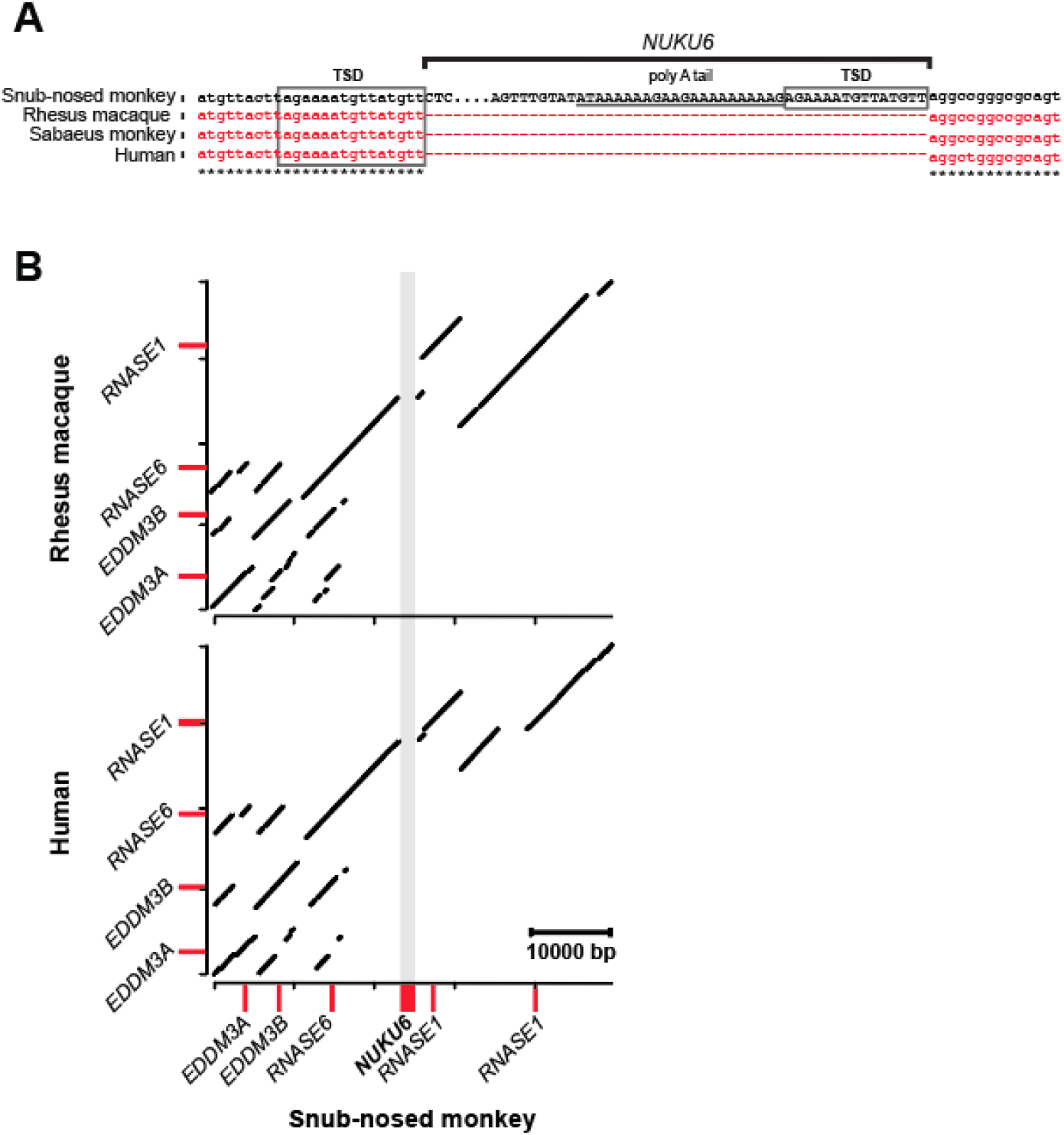
The unique insertion of *NUKU6* into the genome of the golden snub-nosed monkey. (A) Insertion of *NUKU6* compared to the syntenic locus of other primates and evidence of LINE-1 mediated target-site duplication (TSD) and the remnants of an mRNA-derived poly(A) tail. (B) Dot plot representation of the unique insertion of *NUKU6* in the golden snub-nosed monkey genome compared to the syntenic genome position in the human and rhesus macaque genomes.

**Figure S4.**
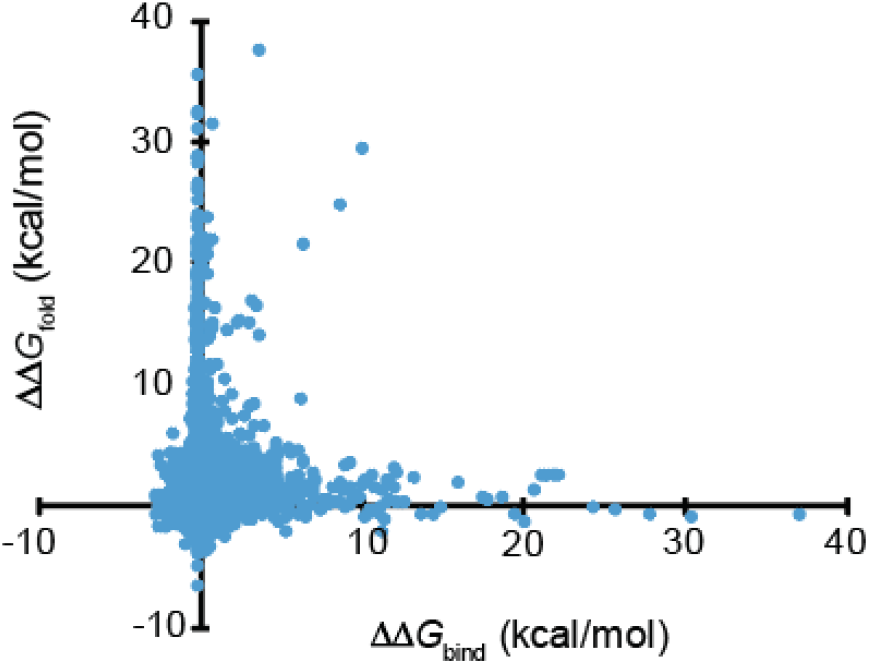
Molecular modeling of the interaction of Ku70p with Ku80p. For Ku70p every possible non-synonymous mutation was plotted to illustrate effects on ΔΔ*G*_fold_. ΔΔ*G*_bind_ was also calculated to predict mutations that would disrupt Ku70p-Ku80p binding.

**Figure S5.**
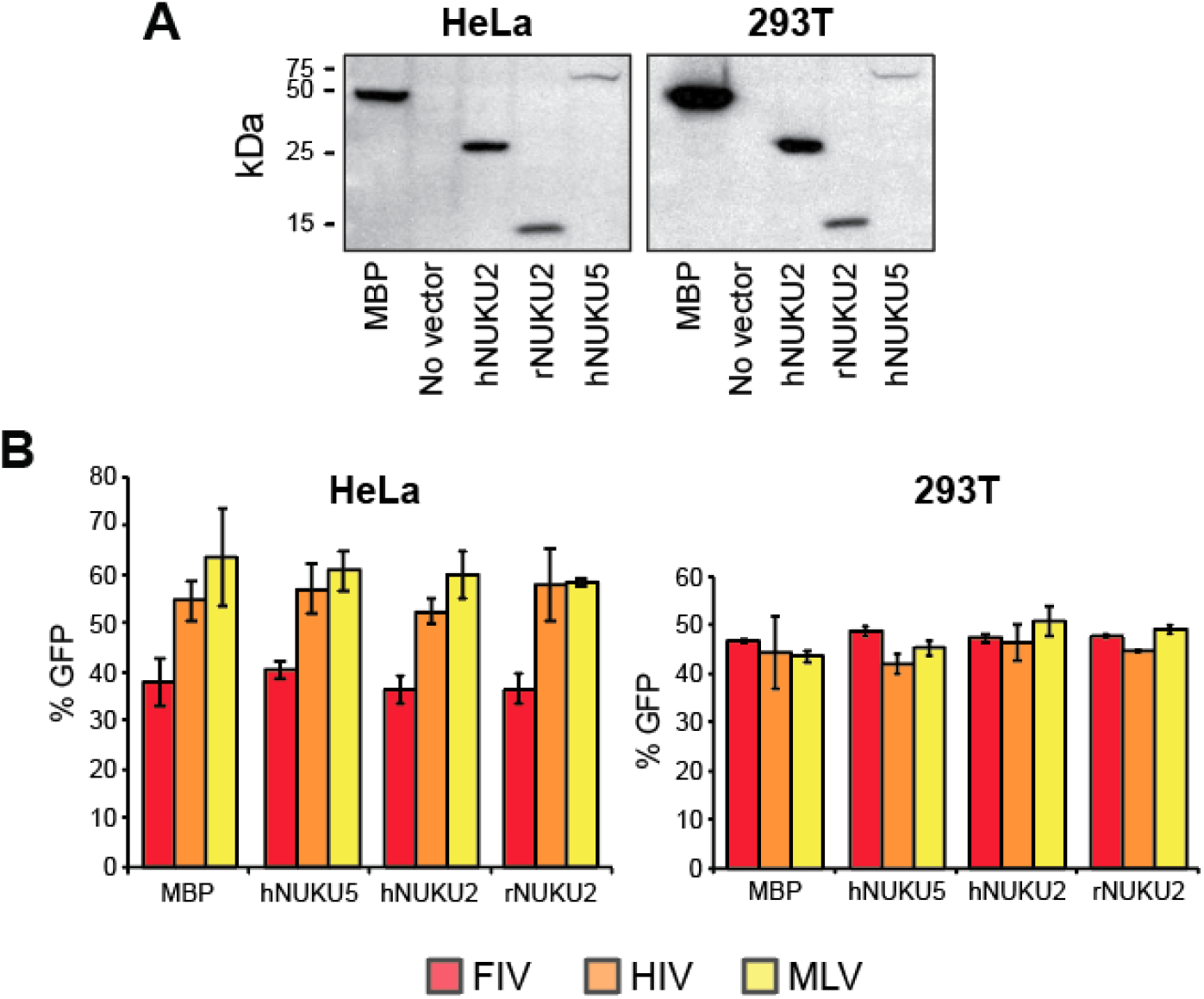
The expression of *NUKU* genes in cell culture does not inhibit retrovirus transduction. (A) The detection by Western blotting of NUKU proteins when transiently expressed in mammalian cell culture. (B) The detection of retrovirus transduction by flow cytometry indicating the percentage of cells that were positive for GFP.

## Supplementary Files

**File S1**. 5’ and 3’ RACE of an unspliced transcript of *NUKU2* from total RNA isolated from human testis.

**Files S2**. A comprehensive list of the change in ΔΔ*G*_fold_ of Ku70p and ΔΔ*G*_bind_ between Ku70p and Ku80p as a result of non-synonymous substitutions in Ku70p.

**File S3**. All nucleotide sequence data for primate *KU70* and *NUKU* retrogenes.

**TABLE S1.** PRIMATE SAMPLES.

| **TABLE S1** | **PRIMATE SAMPLES** |  |  |  |
| --- | --- | --- | --- | --- |
| **Common Name** | **Scientific Name** | **Source** |  | **Cell Type** |
| Gorilla | *Gorilla gorilla* | Coriell | PR00280 | Fibroblasts |
| Sumatran Orangutan | *Pongo pygmaeus* | Coriell | PR01052 | B-Lymphocyte |
| Borneo Orangutan | *Pongo pygmaeus* | Coriell | PR00650 | B-Lymphocyte |
| Siamang | *Hylobates syndactylus* | Coriell | PR00722 | Fibroblasts |
| White-Cheeked Gibbon | *Hylobates leucogeny* | Coriell | PR01037 | Fibroblasts |
| Agile Gibbon | *Hylobates agilis* | Coriell | PR00773 | Fibroblasts |
| Rhesus | *Macaca mulatta* | W. Johnson | Mm265-95 | B-Lymphocyte |
| Talapoin | *Miopithecus talapoin* | Coriell | PR00716 | Fibroblasts |
| Colobus | *Colobus guereza* | Coriell | PR00980 | Fibroblasts |
| Leaf Monkey | *Trachypithecus francoisi* | Coriell | PR01099 | Fibroblasts |
| Crab-eating Macaque | *Macaca fascicularis* | Coriell | Mf103-06 | B-Lymphocyte |
| Olive Baboon | *Papio anubis* | Coriell | PR00978 | Fibroblasts |
| Black Mangabey | *Lophocebus albigena* | Coriell | PR01215 | Fibroblasts |
| Wolf's Guenon | *Cercopithecus wolfi* | Coriell | PR01241 | Fibroblasts |

**Supporting Table S2.**
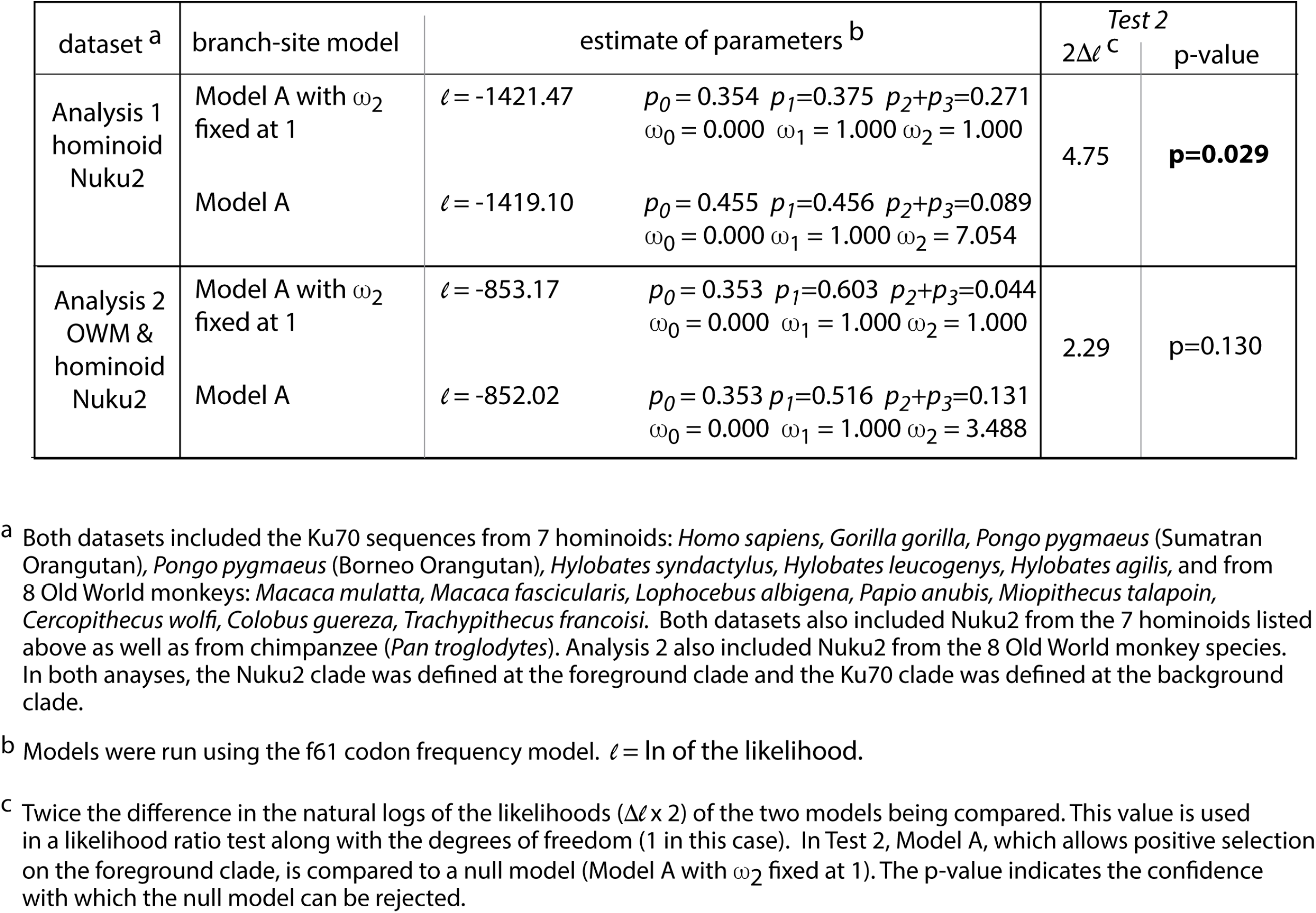
Branch-site test for positive selection of Nuku2.

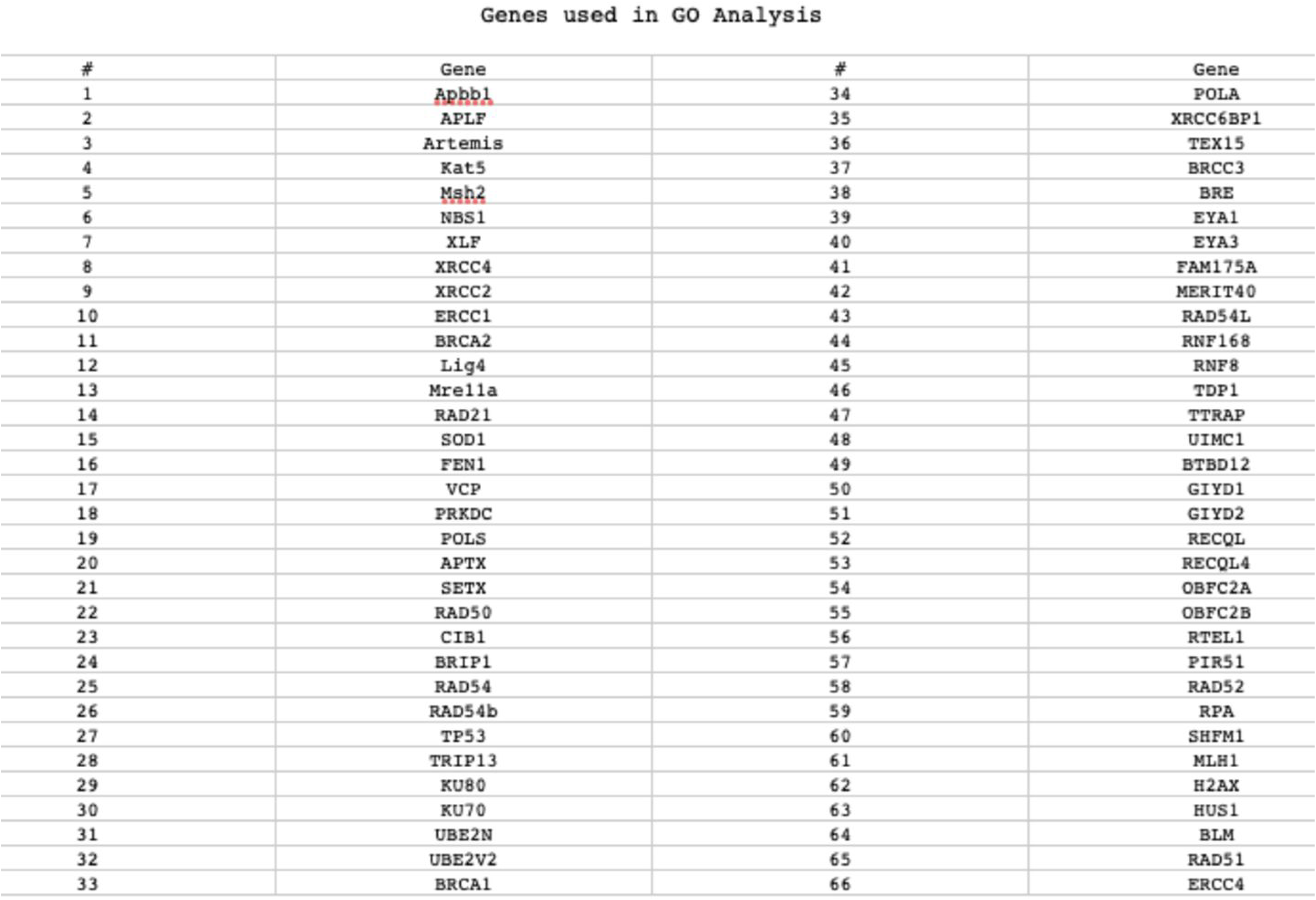

**TABLE S4.** PCR AND SEQUENCING STRATEGIES.

| **TABLE S4** | **PCR AND SEQUENCING STRATEGIES** |
| --- | --- |
| This table lists PCR and sequencing primers used to amplify and sequence Ku70 and Nuku from aforementioned primate samples. |  |
| (PCR primer), *sequencing primer |  |
| **Ku70** |  |
| **ORGANISM** | **PRIMERS** |
| Gorilla PR00280 | (Ai009/Ai016) Ai009*, Ai013*, Ai019*, Ai018*, Ai037*, Ai021*, Ai015*, Ai022a* |
| Borneo Orangutan PR00650 | (Ai009/Ai016) Ai009*, Ai016*, Ai013*, Ai019*, Ai018*, Ai037*, Ai021*, Ai015*, Ai022a* |
| Siamang PR00722 | (Ai009/Ai016) Ai009*, Ai013*, Ai019*, Ai018*, Ai021*, Ai015*, Ai022a* |
| White-Cheeked Gibbon PR01037 | (Ai009/Ai016) Ai009*, Ai013*, Ai019*, Ai018*, Ai037*, Ai021*, Ai015*, Ai022a* |
| Agile Gibbon PR00773 | (Ai009/Ai016) Ai009*, Ai013*, Ai019*, Ai018*, Ai037*, Ai015*, Ai022a* |
| Rhesus Mm265-95 | (Ai009/Ai016) Ai013*, Ai019*, Ai018*, Ai021*, Ai037*, Ai015*, Ai022a* |
| Talapoin PR00716 | (Ai009/Ai016) Ai009*, Ai013*, Ai019*, Ai018*, Ai015*, Ai022a*, Ai037* |
| Colobus PR00980 | (Ai009/Ai016) Ai009*, Ai013*, Ai019*, Ai018*, Ai015*, Ai022a* |
| Leaf Monkey PR01099 | (Ai009/Ai016) Ai009*, Ai013*, Ai018*, Ai015* |
| Crab-eating Macaque Mf103-06 | (Ai009/Ai016) Ai009*, Ai013*, Ai019*, Ai018*, Ai037*, Ai015* |
| Olive Baboon PR00978 | (Ai009/Ai016) Ai009*, Ai016*, Ai014*, Ai013*, Ai019*, Ai037*, Ai021*, Ai022a* |
| Black Mangabey PR01215 | (Ai009/Ai016) Ai004*, Ai013*, Ai015*, Ai019*, Ai037*, Ai022a* |
| Wolf's Guenon PR01241 | (Ai009/Ai016) Ai013*, Ai019*, Ai009*, Ai018*, Ai037*, Ai015*, Ai021*, Ai022a* |
| **Nuku2** |  |
| **ORGANISM** | **PRIMERS** |
| Gorilla PR00280 | (Ai023/Ai028) Ai038*, Ai023*, Ai054*, Ai024*, Ai025*, Ai028* |
| Borneo Orangutan PR00650 | (Ai023/Ai028) Ai038*, Ai023*, Ai054*, Ai024*, Ai025*, Ai028* |
| Siamang PR00722 | (Ai022b/Ai030) Ai038*, Ai023*, Ai054*, Ai024*, Ai025* |
| White-Cheeked Gibbon PR01037 | (Ai023/Ai028) Ai038*, Ai023*, Ai054*, Ai053*, Ai025*, Ai055* |
| Agile Gibbon PR00773 | (Ai023/Ai028) Ai038*, Ai023*, Ai054*, Ai024*, Ai053*, Ai055*, Ai025* |
| Talapoin PR00716 | (Ai023/Ai028) Ai038*, Ai023*, Ai024*, Ai028*, Ai025* |
| Colobus PR00980 | (Ai023/Ai028) Ai038*, Ai023*, Ai024*, Ai026*, Ai028*, Ai025* |
| Leaf Monkey PR01099 | (Ai023/Ai028) Ai038*, Ai023*, Ai024*, Ai028*, Ai025* |
| Crab-eating Macaque Mf103-06 | (Ai023/Ai028) Ai038*, Ai023*, Ai024*, Ai028*, Ai025* |
| Olive Baboon PR00978 | (Ai023/Ai028) Ai038*, Ai023*, Ai054*, Ai024*, Ai055*, Ai025*, Ai028* |
| Black Mangabey PR01215 | (Ai023/Ai028) Ai038*, Ai023*, Ai054*, Ai024*, Ai055*, Ai025*, Ai028* |
| Wolf's Guenon PR01241 | (Ai023/Ai028) Ai038*, Ai023*, Ai054*, Ai024*, Ai055*, Ai025*, Ai028* |

**TABLE S5.**
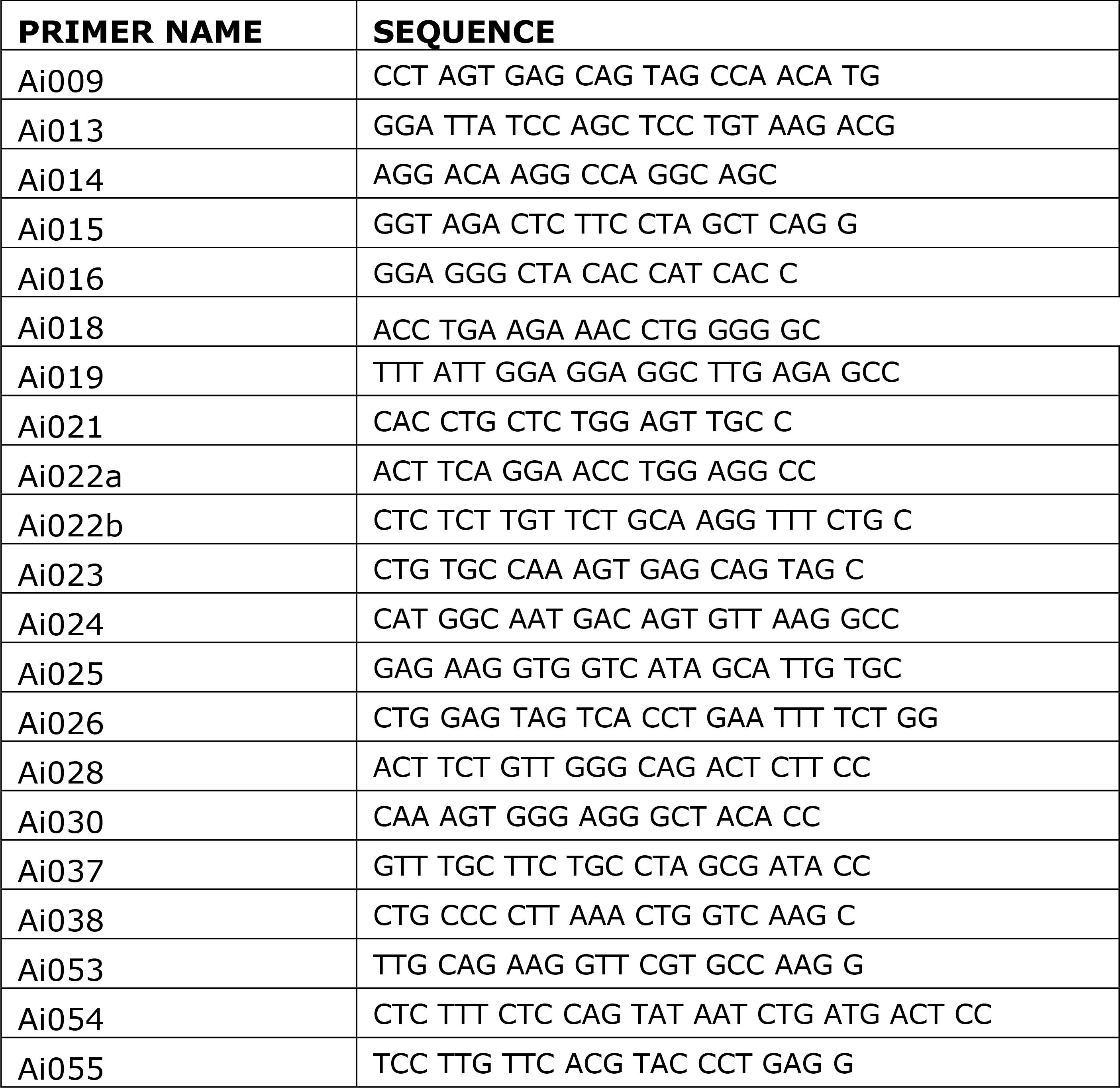
PRIMERS USED FOR AMPLIFICATION AND SEQUENCING OF KU70 AND NUKU2.

